# Open-source 3D printed manifolds for exposure studies using human airway epithelial cells

**DOI:** 10.1101/2024.08.12.607646

**Authors:** Ryan Singer, Elizabeth Ball, Nadia Milad, Jenny P. Nguyen, Quynh Cao, Ravi Selvaganapathy, Boyang Zhang, Mohammadhossein Dabaghi, Imran Satia, Jeremy A. Hirota

## Abstract

**Rationale:** Inhalation of airborne stimuli can damage the airway epithelium, increasing the risk of developing respiratory or systemic diseases. *In vitro* studies using air-liquid interface cell cultures enable controlled investigation of cellular responses to relevant exposures. Commercial *in vitro* exposure systems provide precise and reproducible dosage but require significant capital investment and are not amenable to customization. Research groups interested in respiratory exposure science may benefit from a more accessible alternative open-source exposure system. We present 3D printed manifolds for applying a range of airborne exposures uniformly across standard, commercially available 6- and 24-well plates with air-liquid interface culture inserts.

**Methods:** A simple chamber-style exposure system and the manifolds were evaluated for exposure uniformity via computational fluid dynamics simulations and deposition of nebulized FITC-labelled dextran. The chamber and manifolds were manufactured using a stereolithography 3D printer. Cannabis concentrate vapor was generated from 3 different vaporizers and applied to well plates using the manifold system. Calu-3 cells were cultured on Transwell™ inserts and exposed to whole tobacco smoke or room air.

**Results:** The manifolds produced less variation in simulated air velocities and physical deposition of FITC-dextran aerosol deposition across well plates compared to those of the chamber-style exposure system. Distinct doses of cannabis concentrate vapour were delivered to well plates with low variation among wells. Whole tobacco smoke exposure using the manifold system induced functional changes in Calu-3 airway epithelial cell barrier function, cytokine production (IL-6 and IL-8), and cell membrane potential.

**Conclusions:** Collectively, our data demonstrate the feasibility and the validity of our open-source 3D printed manifolds for use in studying various respiratory exposures and position our designs as more accessible options in parallel with commercially available systems.

All article content is licensed under a Creative Commons Attribution (CC BY-NC 4.0) license (https://creativecommons.org/licenses/by-nc/4.0/).

## Introduction

Breathing results in exposure to potentially noxious airborne pathogens, irritants, and gases which could influence the development and progression of lung diseases. The first point of contact for inhaled stimuli is the respiratory airway epithelium, which provides protective mechanisms including mucus secretion and tight junctions to form a physical barrier that is paired with production of soluble immune mediators and antimicrobial proteins [1–3]. Despite this tiered layering of protection, repeated exposure to external stimuli can damage the airway epithelium, increasing the risk of developing airway diseases including asthma, COPD, and lung cancer [4, 5].

Controlled environmental exposure studies can be conducted in humans, *in vivo* animal models, and *in vitro* with diverse lung cell types [6–10]. *In vitro* systems provide high experimental control but often use epithelial cells cultured in submerged monolayers, which lack the complexity of real-world airborne exposures and certain *in situ* cellular characteristics [11, 12]. Advanced models use airway epithelial cells at an air-liquid interface (ALI), promoting cell differentiation and development of *in situ*-like properties and transcriptomics [13–15]. Commercial systems like the Cultex® Radial Flow, Vitrocell® Continuous Flow and Cloud Alpha, and Scireq® ExpoCube® enable direct ALI exposures [16–19]. These systems provide precise dosimetry but require significant investment and are not amenable to customization for diverse applications. In-house bespoke systems offer alternatives but often lack uniformity, throughput, or reproducibility [10, 20, 21]. The simplest custom approach for applying *in vitro* environmental exposures is a chamber-style system in which exposures are applied to a common volume containing the cell cultures. Such systems may lack exposure uniformity and efficiency, with only a small portion of the stimulus contacting the cells [19]. The disadvantages of current *in vitro* respiratory exposure systems suggest that there is an unmet need for systems that are accessible, adaptable, and most importantly, reproducible.

Respiratory exposure science research may benefit from an open-source low-cost alternative exposure system for performing exploratory *in vitro* environmental exposures. In this study, we present 3D printed manifolds for applying a range of airborne exposures uniformly across standard, commercially available well plates with ALI culture inserts. The aim of this study was to validate the utility of the 3D printed manifolds for delivering nebulized aerosols and cannabis concentrate vapour to standard well plates and for investigating *in vitro* responses of airway epithelial cells to tobacco smoke.

## Methods

### Computational fluid dynamics (CFD)

3D models of the manifolds mounted to 6- and 24-well plates with Transwell™ inserts were designed using Autodesk® Fusion 360 (v16.4). Standard ACIS Text (.sat) files were exported to Autodesk® CFD 2023, and the Geometry Tools function was used to generate internal volumes defining the air space within the manifold-plate systems. These internal volumes were assigned the fluid properties of air as defined by the default material library. The following CFD decisions followed best practices from a recent review on computational modeling of fluids [22]. Static non-slip conditions were applied to all outer boundaries except the outlets, which were open to atmosphere, and the inlet, which was defined as 2.5 and 10 mL/s flow rate for the 6- and 24-well manifolds, respectively. Meshing was automatically generated with surface and gap refinement disabled. The meshes were inspected visually before computation to ensure regions of interest were adequately discretized. The air volumes were assumed as incompressible with a k-epsilon turbulence model. Solutions were computed until convergence with Intelligent Solution Control enabled and Automatic Convergence Assessment set to default. Mean static pressure, maximum velocity, and maximum shear stress were quantified via profile planes generated at the approximate cell culture surface, from which the coefficients of variation (CV = standard deviation/mean) were calculated among the wells.

### Manifold fabrication

The manifolds were designed using Autodesk® Fusion 360 with reference to caliper measurements and technical data sheets of Corning™ Costar™ and Eppendorf® well plates. Stereolithography files were exported to PreForm 3.37.3 software (Formlabs) with the manifold oriented upright. Support density and touchpoint size were set to 0.9 and 0.4, respectively, and the part was sliced with a resolution of 0.1 mm. A transparent resin (Clear V4, Formlabs, product No. RS-F2-GPCL-04) was used to print the manifolds with an SLA resin printer (Form 3B, Formlabs, product No. PKG-F3B-WS-SVC). **Supplementary Figure 1** illustrates the post-print processing. All manifold and base design files are provided in the Online Supplement in stereolithography (.STL) and Standard ACIS Text (.SAT) formats and available for use under Creative Commons licensing (CC BY-NC 4.0).

### Aerosol distribution

A micropump nebulizer (Aeroneb® Lab, Kent Scientific Corp., product No. NEB7000 and NEB1000) was used to generate an aerosol from 2 mg/mL 4kDa FITC-dextran (Chondrex Inc., product No. 4013) dissolved in deionized water. The aerosol was accelerated using a 50 mL syringe connected to the nebulizer base inlet. The outlet was connected via 5 cm long, 8 mm inner diameter E-3603 Tygon tubing (McMaster Carr, product No. 5186T16) to manifolds mounted on well plates (**Supplementary Figure 2A**). One puff of aerosol was defined as a 5 second plunge of the syringe at a rate of 10 mL/s for the 24-well manifold and 2.5 mL/s for the 6-well manifold and for the cloud chamber. The manifold was removed for 30 seconds between each puff to prevent stagnation of exposures. Ten fluorescence microscope images (EVOS M7000) at 10× magnification were acquired from each well before exposure and after 1, 3, 6, and 9 puffs of aerosol. Statistical differences in aerosol deposition between puffs were evaluated via one-way ANOVA with Tukey’s post-hoc test. The mean deposition area of the aerosol in each well was quantified using ImageJ. The experiment was repeated using a cloud chamber exposure system to compare the mean variation coefficient (SD/mean) of aerosol deposition to that of the manifolds.

### Cannabis concentrate vapour exposure

Cannabis concentrate purchased from the Ontario Cannabis Store (Cherry Blossom, Growtown Inc., CBD 759mg/g, THC 31 mg/g, CBG 1mg/g) was vaporized using 3 separate rechargeable vaporizer pens: Pen A - Feather (GTIN: 628110247711); Pen B - Good Supply (GTIN: 694144004354); and Pen C - Yocan Kodo Pro (Shenzhen, China, unknown product number). The voltage was varied between low (2.8 V) and high (3.6 V) settings using the Good Supply and Kodo Pro Yocan vaporizers, whereas the Feather vaporizer operated at a fixed voltage. Each vaporizer was used to administer 9 puffs of vaporized cannabis concentrate onto cell-free 24-well plates using the 24-well manifold with all wells containing 300 µL of Minimum Essential Medium Alpha. One puff was defined as 50 mL of smoke or air perfused through the 24-well manifold at 10 mL/s using a 3-way valve connected to the manifold, a 50 mL syringe, and a vaporizer. Media was subsequently collected and stored at -80 until cannabidiol (CBD) concentrations were measured via mass spectrometry.

### Mass spectrometry measurement of CBD

Sample aliquots were thawed on ice and mixed with 400 μL of cold acetonitrile containing 1% (v/v) formic acid and 50 ng/mL of internal standard mixture. Samples were vortexed for 1 minute followed by centrifugation at 18,000 x g for 15 minutes. A 100 μL aliquot of the top organic layer was transferred to an HPLC vial with glass insert and combined with 100 μL LCMS grade water. The tube was vortexed for 10 seconds prior to injection of 5 μL into an Agilent 1290 Infinity II HPLC coupled to an Agilent 6495C iFunnel QQQ mass spectrometer (Agilent, Santa Clara, CA, USA). Certified CBD standards (Supelco, product No. C-045) and corresponding deuterated internal standards (Cerilliant, product No. C-084) were used to quantify the amounts in the media samples. The internal standards were added to all calibration standards, test samples, and quality control (QC) samples. Statistical differences between groups were evaluated via one-way ANOVA with Tukey’s post-hoc test.

### Cell culture

The Calu-3 epithelial cell line was cultured in Transwell™ inserts as previously described [11, 23]. Briefly, Calu-3 cells were expanded in tissue culture flasks before seeding into Transwell™ inserts (seeding density 150,000 cells/insert) in a 24-well plate. Cells were fed every 2 days with Minimum Essential Medium Alpha (Corning, product No. CA45000-300) supplemented with 10% fetal bovine serum (FBS; Wisent Inc., product No. 080-450), 1% antibiotic/antimycotic (Thermo Fisher, product No. 15240062), and 1% HEPES buffer (Corning, product No. 25-060-CI). When cells were confluent, they were subjected to ALI by removing the apical media to ensure exposures directly contacted cells. Exposures were commenced on day 0 of ALI conditions.

### Tobacco smoke experiment

#### Exposure

Calu-3 cells on Transwell™ inserts were exposed to 3 or 6 puffs of tobacco smoke (3R4F, University of Kentucky) or 6 puffs of room air, where one puff consisted of 50 mL of smoke or air perfused through the 24-well manifold at 10 mL/s. Exposures were generated using a 3-way valve connected to the manifold, a 50 mL syringe, and a cigarette or room air (**Supplementary Figure 2B**). The manifold was removed for 30 seconds between each puff to purge the previous puff and prevent stacking of exposures.

#### Outcomes

Brightfield images (EVOS M7000) and trans-epithelial electrical resistance (TEER; an indicator of epithelial barrier integrity) measurements (Millicell ERS-2) were gathered and calculated pre-exposure and 18 h post-exposure. Basal media was collected 18 h post-exposure for analysis of lactate dehydrogenase (LDH), IL-6, and IL-8 concentrations. LDH was measured using the CyQUANT™ LDH Cytotoxicity Assay (Invitrogen, product No. C20301) as a marker of cell death. Pro-inflammatory cytokines IL-6 and IL-8 were measured using DuoSet ELISA kits (R&D Systems, product No. DY206 and DY208) following manufacturer’s instructions. Cell membrane potential, a marker of ion channel activity, was measured via fluorescence-indicating dye 20 h post-exposure using the SpectraMax® i3x plate reader (Molecular Devices, Product No. i3x) and the FLIPR® membrane potential assay kit (Molecular Devices, Product No. R8042), following manufacturer’s instructions and using methods previously described [12, 24]. Briefly, cells were washed with HBSS before loading the dye-containing solution, and the cell culture plate was then placed into the plate reader. Baseline fluorescence was recorded for 40 min before the addition of 10 µM forskolin to half of the samples to stimulate ion channel activity, while the remaining samples received a vehicle for unstimulated controls. Fluorescence was measured 6 times at 5-minute intervals following the addition of forskolin or vehicle. Measurements from forskolin-stimulated samples were normalized to the unstimulated room air-exposed samples, and the maximum peak and area under curve (AUC) of normalized fluorescence intensity were determined. Statistical differences between groups were evaluated via t-tests.

## Results

### Chamber-style system fluid dynamics simulation and physical aerosol deposition

The simplest exposure system that can be fabricated with 3D printing is a chamber box with an inlet for applying exposures (**Figure 1A**). The effectiveness of this simple design was interrogated with fluid dynamic simulations and lab experiments with FITC-labelled dextran nebulized into the chamber.

**Figure 1.**
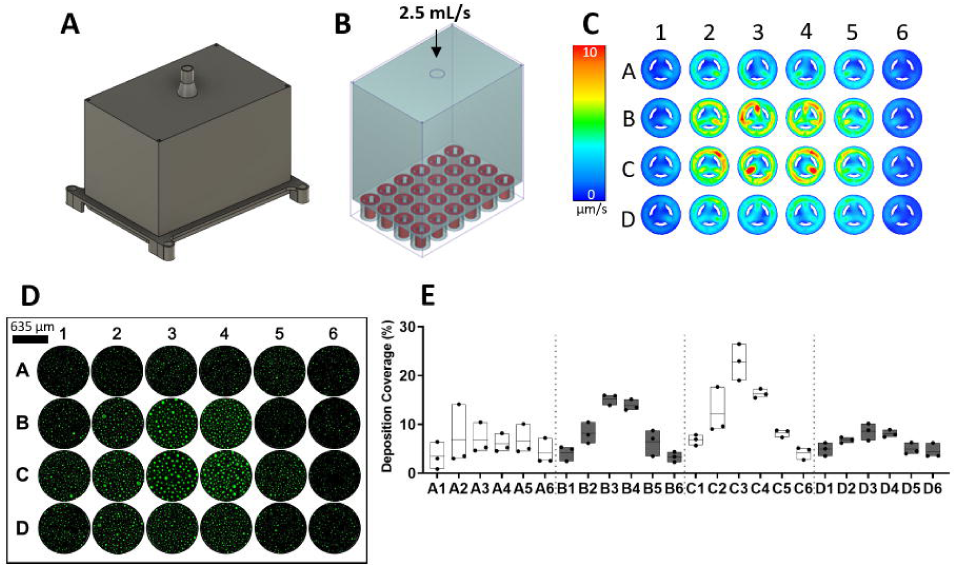
Distribution of air and nebulized FITC-dextran in a simple chamber-style exposure system. A fluid dynamics simulation and perfusion of an aerosolized fluorescent solution were performed. Renderings of **A)** the chamber-style well plate lid and **B)** the air space (blue) within the chamber-well plate system with modelled Transwell™ inserts (red). **C)** Planar velocity profile at the surface of the Transwell™ inserts. **D)** Representative microscope images (10×) and **C)** quantified deposition coverage of FITC-dextran aerosol droplets in each well of a 24-well plate where each bar spans the minimum to maximum value with a mean centre line.

Using a fixed inlet air flow of 2.5 mL/s through a 6.5 mm diameter designed inlet (**Figure 1B**), we performed a steady state simulation of air flow in the chamber-style exposure system, revealing that the planar velocity profile at the surface of the Transwell™ inserts displayed maximum flow velocities ranging from 16 µm/s in a centre well (B4) to 2.7 µm/s in a peripheral well (D6) of the 24-well plate (**Figure 1C**). The mean flow velocity at the surface of the Transwell™ inserts across all wells was 1.46 µm/s with standard deviation of 0.93 µm/s, resulting in a coefficient of variation of 63.7%. Within the chamber, the highest flow velocity was 5.89 cm/s immediately beneath the inlet in the centre of the well plate with a gradient of decreasing flow velocity toward the periphery of the plate. This analysis was carried out with multiple fluid dynamic simulation conditions with varying inlet diameters and flow rates, yielding similar variability of air flow rates at the surface of the Transwell™ inserts. These results suggest that physical aerosol deposition in a chamber-style system would be highly variable within a well plate.

To functionally test the deposition within a chamber-style system, aerosol exposures of FITC-labelled dextran (2 mg/mL) were applied to 3D printed chambers mounted on 24-well plates at a flow rate of 2.5 mL/s. The deposition of the FITC-labelled dextran was quantified in each well. The percentage area of deposited aerosol droplets was highest in the centre wells (well locations B3, B4, C3, and C4) compared to peripheral wells (**Figure 1D-E**). The mean area of a given well covered by FITC-dextran was 7.41% with a standard deviation of 5.32%, resulting in a 71.8% coefficient of variation across the wells.

Our modeling and functional assessment of a simple chamber-style design suggest that a high degree of variability of deposition would result across a standard 24-well plate and may introduce undesired heterogeneity into exposure science experiments.

### Manifold system fluid dynamics simulation and physical aerosol deposition

To reduce or eliminate observed heterogeneities in deposition we designed manifold systems that included a central chamber with distribution channels designed to reduce heterogeneity across the wells of 6- and 24-well plates. As with the chamber-style system, computational fluid dynamic simulations were performed in advance of 3D printing to refine manifold specifications prior to functional testing.

Models of a 6- and 24-well manifold system were developed in Autodesk (see methods), with design constraints for accommodating the dimensions of common commercially available well plates (**Figure 2A, E**). The final models were used for the computational fluid dynamics simulation (**Figure 2**) with prior design iterations and simulations excluded from the present analysis. Design optimizations included changing channel length, diameter, and contour in addition to changing the volume of the chamber feeding the channels.

**Figure 2.**
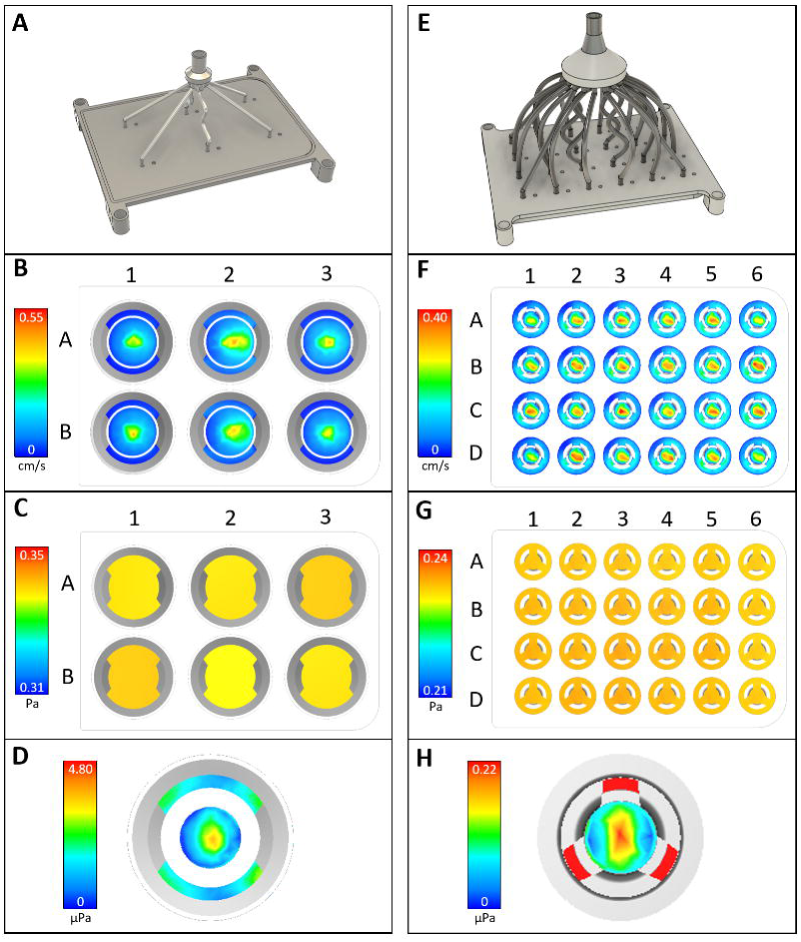
Fluid dynamics simulations of air flow in well plates using the 6-well and 24-well manifolds. Renderings of the **A)** 6-well and **E)** 24-well manifolds. Planar velocity profiles at the Transwell™ membrane surface in each well for the **B)** 6 well and **F)** 24-well manifolds. Planar pressure profiles near the inlet of the manifold channels in each well for the **C)** 6-well and **G)** 24-well manifolds. Representative shear stress profiles near the Transwell™ membrane surface for the **D)** 6-well and **H)** 24-well manifolds.

For the optimized manifold designs, air flow simulations demonstrated uniformity in velocity profiles at the surface of the Transwell™ inserts across all wells using the 6-well (0.56 ± 0.06 cm/s) and 24-well (0.40 ± 0.05 cm/s) manifolds with coefficients of variation of 11.3% and 11.8% respectively (**Figure 2B, F**). The mean static pressure among all wells of 6-well (0.247 ± 0.003 Pa) and 24-well (0.233 ± 0.001 Pa) plates resulted in coefficients of variation of 1.0% and 0.3% respectively (**Figure 2C, G**). Shear stress at the cell culture surface was 4.27 ± 0.356 µPa and 0.157 ± 0.014 µPa for the 6- and 24-well manifolds, respectively, resulting in coefficient of variation of 8.3% and 8.9% (**Figure 2D, H**).

The computational fluid dynamics data suggest that the manifold designs reduce heterogeneity of exposure across a standard well plates compared to a simple chamber-style system. We therefore completed design and 3D printed fabrication of the 6- and 24-well manifold configurations (**Figure 3**). The final design for both plate configurations included a base plate that was printed to hold the standard well plate. Assembly of the system requires enclosing the well plate between the base and the manifold using the magnets to secure, ensuring all edges of the well plate are contacting the gasket.

**Figure 3.**
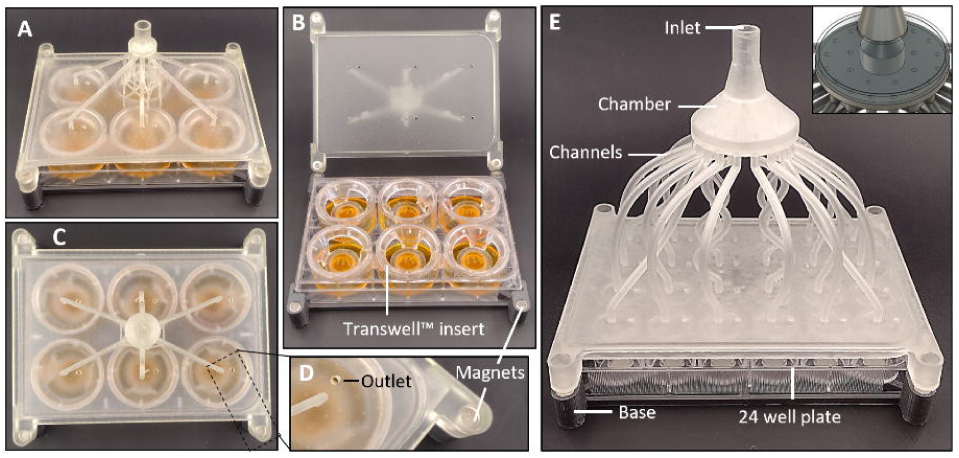
Structure of the 3D printed manifolds. Photos of the 6 well manifold system in perspective view with the manifold **A)** closed and **B)** open. **C)** Top view photo of the 6 well manifold system with **D)** close-up on an outlet and a magnet slot. **E)** Perspective view photo of the 24 well manifold system.

Following fabrication, functional testing was performed using aerosolization of FITC-dextran as described above, with an additional concentration-response component added to this characterization (**Figure 4**). The percentage area of aerosol deposition was proportional to the number of puffs for both the 6- and 24-well manifolds, with significantly different amounts of deposition with 1, 3, 6, and 9 puffs (**Figure 4A-B**). The mean percentage area of aerosol deposition following 9 puffs was not significantly different across the wells of 6-well (43.57 ± 6.41%) and 24-well (22.38 ± 2.61%) plates, with coefficients of variation of 14.7% and 11.7% respectively (**Figure 4C-F**).

**Figure 4.**
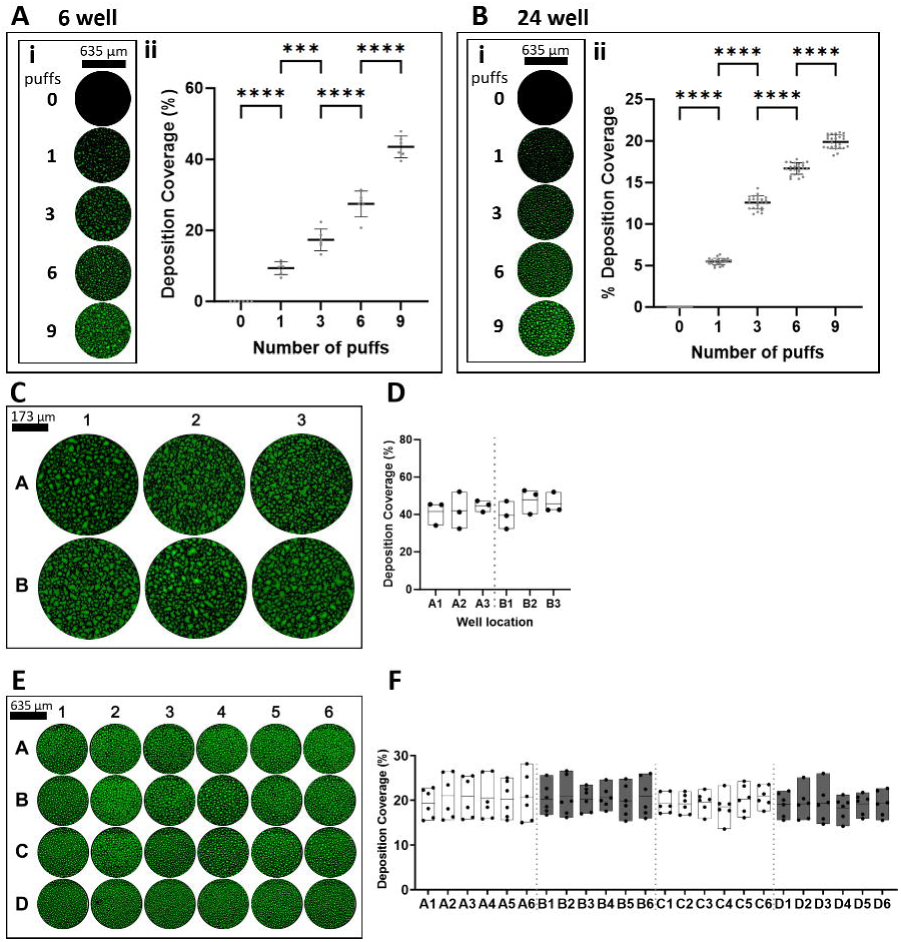
Nebulized FITC-dextran aerosol deposition in the 6-well and 24-well manifold systems. Deposition with increasing number of puffs using the **A)** 6-well and **B)** 24-well manifolds with **i)** representative microscope images (10×) and **ii)** quantified deposition coverage of FITC-dextran aerosol droplets with mean and standard deviation. Representative microscope images (10×) of FITC-dextran aerosol deposition across **C)** 6-well and **E)** 24-well plates with **D,F)** quantified deposition coverage in each well where each bar spans the minimum to maximum value with a mean centre line. *** = P ≤ 0.001, **** = P ≤ 0.0001.

### Cannabis concentrate vapour distribution with commercially available vaping devices

The computational modeling and functional testing of the manifold system suggested the design could have value for real-world respiratory exposure experiments with clinically relevant reagents. The impact of vaporized cannabis concentrates, a combustion-free method of cannabis use, on lung cell biology is only beginning to be explored and may be an application for our manifolds. We therefore explored the ability of the manifold design to deliver cannabis concentrate aerosols generated from three commercially available vaping devices. We chose three devices with different features including breath-activated heating, button-activated heating, and the ability to control the voltage delivered to the heating coil that generates the vapour (**Figure 5A**). For each of the vaping devices we used the same cannabis concentrate solution with high CBD and low THC (Cherry Blossom, Growtown Inc., CBD 759mg/g, THC 31 mg/g, CBG 1mg/g). The cannabis concentrate was delivered to well plates via 9 puffs of vapour using the 24-well manifold. The breath-activated Pen A delivered the lowest dose of 1.97 ± 1.44 µg/mL CBD, having a 73% coefficient of variation. Pen B delivered 14.0 ± 0.81 µg/mL CBD with low voltage (2.8 V) and 16.1 ± 0.55 µg/mL CBD with high voltage (3.6 V), yielding coefficients of variation of 5.8% and 3.4% respectively. Pen C delivered 22.4 ± 0.58 µg/mL CBD with low voltage and 25.9 ± 1.13 µg/mL CBD with high voltage, yielding coefficients of variation of 2.6% and 4.4% respectively (**Figure 5B**). Our results indicate that our manifold designs are amenable to cannabis concentrate vapour exposure experiments and can be used to validate the reproducibility/consistency of cannabis delivery with different vape pens.

**Figure 5.**
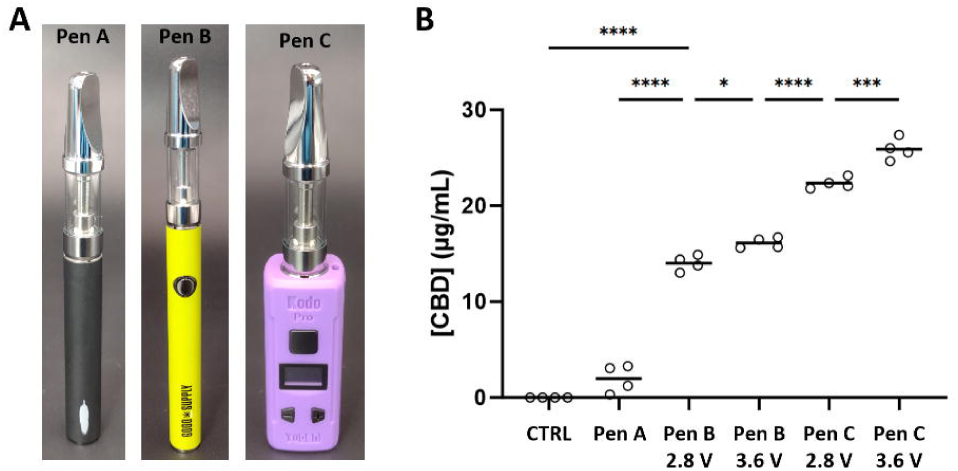
Application of distinct cannabidiol doses using different cannabis concentrate vaporizers. **A)** Photos of each cannabis concentrate vaporizer, “Pens A-C”. **B)** Cannabidiol (CBD) concentration delivered to well plates via the 24-well manifold using each vaporizer. Pen A is breath-activated, and Pens B and C have adjustable heating coil voltage between low (2.8 V) and high (3.6 V) settings. * = P ≤ 0.05, *** = P ≤ 0.001, **** = P ≤ 0.0001.

### Functional consequences of tobacco smoke exposure on Calu-3 human airway epithelial cells

Tobacco smoke exposure can impact human airway epithelial cell phenotype and function with changes in cell viability, cytokine production, barrier function, and transport mechanism [11, 23]. We therefore opted to validate the manifold designs with the Calu-3 human airway epithelial cell line with 6 puffs of whole tobacco smoke and 6 puffs of room air control exposures. Brightfield images pre-exposure were similar across all samples, whereas images 18 h post-exposure revealed areas void of cells in tobacco smoke-exposed samples with no apparent change in cell morphology (**Figure 6A**). Room air exposure induced a 10.5 ± 7.5% TEER reduction, whereas tobacco smoke exposure induced a significantly larger TEER reduction of 54.5 ± 10% from the pre-exposure baseline 18 h post-exposure (**Figure 6B**). Mean LDH release into the basolateral compartment of the ALI cultures was not significantly different between room air (5.22 ± 1.54%) and tobacco smoke (9.52 ± 6.68%) exposure conditions 18 h post-exposure (**Figure 6C**). Tobacco smoke exposure induced increases in pro-inflammatory IL-6 (80.97 ± 13.70 pg/mL) and IL-8 (1548 ± 14.05 pg/mL) in the basal cell culture media 18 h post exposure relative to the room air control levels of IL-6 (18.47 ± 3.178 pg/mL) and IL-8 (1401 ± 54.44 pg/mL) (**Figure 6D-E**).

**Figure 6.**
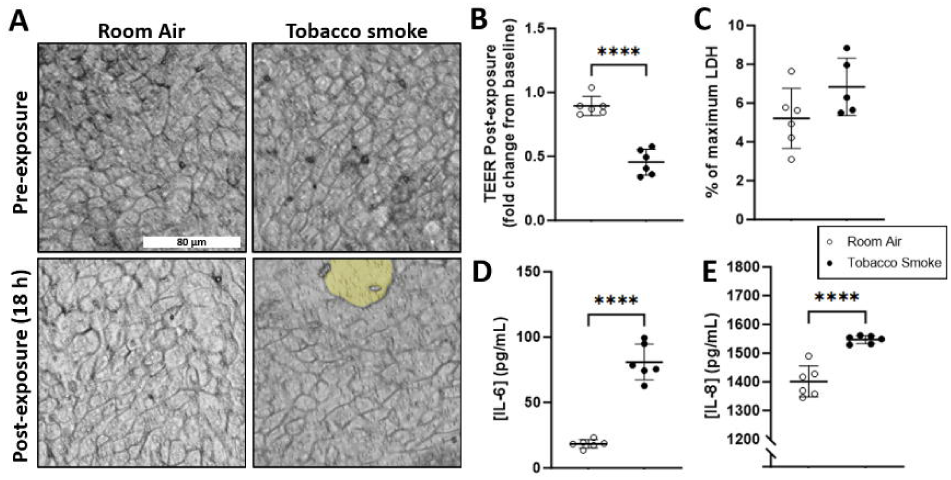
Responses of human airway epithelial cells to whole tobacco smoke exposure. Calu-3 epithelial cells at ALI were exposed to 6 puffs of room air or tobacco smoke using the 24-well manifold. **A)** Representative brightfield images (20×) pre-exposure and 18 h post-exposure. **B)** Trans-epithelial electrical resistance (TEER) 18 h post-exposure relative to pre-exposure measurements. **C)** Lactate dehydrogenase (LDH) relative to maximum release 18 h post-exposure. **D)** IL-6 and **E)** IL-8 concentrations in the basal compartment of ALI cultures 18 h post-exposure. **** = P ≤ 0.0001.

Tobacco smoke exposure has been associated with a down-regulation of CFTR protein and corresponding reduction in membrane transport of chloride and bicarbonate ions [12]. Fluorescence plate-based assays for CFTR function are high-throughput approaches for indirectly measuring ion transport through monitoring of membrane potential change. In these assays, forskolin is used as an adenylyl cyclase activator to increase intracellular cAMP levels, activate PKA, phosphorylate CFTR, and increase channel opening probability and ion flux that can be measured as a change in membrane potential. In room air exposed airway epithelial cells the membrane potential changes induced by forskolin treatment had a maximum peak value of 2.63 ± 0.21 arbitrary units with AUC of 3739.5 ± 323.1 (**Figure 7**). Tobacco smoke resulted in a significant exposure-dependent decrease in maximum peak and AUC values (3 puffs 1.84 ± 0.03 arbitrary units and AUC 2624.7 ± 45.32; 6 puffs 1.15 ± 0.11 arbitrary units and AUC 1648.3 ± 179.6). Collectively, our data supports the use of our manifold for controlled delivery of tobacco smoke to airway epithelial cells on Transwell™ inserts and provide a validation for future exposure experiments with other combusted products (e.g., cannabis smoke or wood smoke exposures).

**Figure 7.**
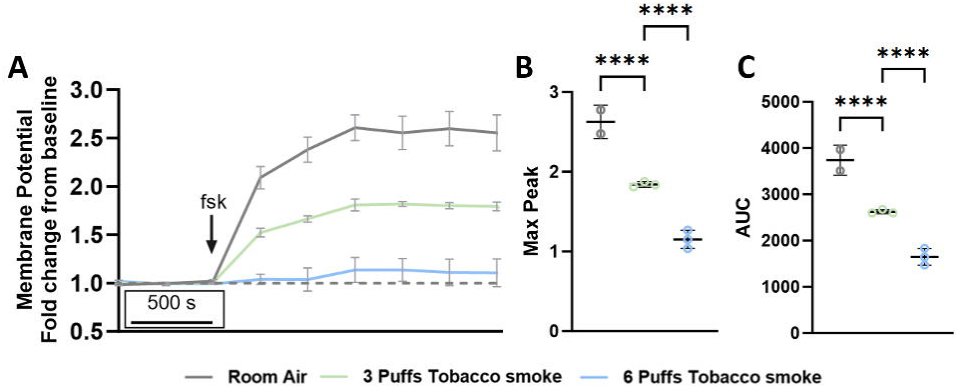
Attenuated cell membrane potential measurements of human airway epithelial cells with forskolin stimulation in response to whole tobacco smoke exposure. **A)** Cell membrane potential relative to non-forskolin-stimulated measurements as a function of time. Addition of forskolin (fsk) is indicated with an arrow. **B)** Maximum peak and **C)** area under curve (AUC) values were interpolated from the time curves. **** = P ≤ 0.0001.

## Discussion

This work presents open-source 3D printed manifolds for applying a range of airborne environmental exposures to standard well plate configurations of ALI cell cultures. We validate the manifolds’ utilities for exposures to 6- and 24-well plates using 1) nebulized aerosols with uniform distribution of FITC-dextran droplets, 2) cannabis concentrate vaping with precise doses of CBD, and 3) tobacco smoke with measurements of exposure-induced changes in Calu-3 epithelial cell TEER, cytokine release, and membrane potential. The 3D printed manifolds provide a rapid and cost-effective method for performing *in vitro* exposure science experiments with broad applications that may include, but are not limited to, gas, aerosol, particulate matter, and smoke exposures.

Many environmental agents impact lung health and are associated with pulmonary and systemic pathologies. Data from the Global Burden of Disease, Injuries, and Risk Factors Study 2019 revealed that lower respiratory tract infections were the leading cause of infectious deaths globally in 2019 [25, 26]. Approximately 16% of deaths per year globally are attributed to pollution, a statistic that has risen 7% since 2015 and 66% since 2000 [27]. Collectively, chronic respiratory diseases were the third leading cause of death globally in 2019, for which the highest risk factor was smoking, followed by air pollution and occupational hazards [4]. Although smoking rates have declined 28% from 1990 to 2019, population growth has led to an increase in the total number of smokers from 0.99 billion to 1.14 billion [28]. The legalization of cannabis in many countries across the globe is also contributing to increased inhaled combustible and aerosol exposures, even though the respiratory effects of these exposures are not well understood [29]. Considering these concerns surrounding environmental exposures and their impact on lung health, the need for robust data on the effects of inhaled stimuli on respiratory pathophysiology is paramount and increasing the accessibility of experimental approaches for *in vitro* exposures will facilitate research and provide alternative methods for use by scientific and regulatory communities.

The leading commercial *in vitro* exposure systems are powerful tools for investigating the responses of lung epithelial cells to direct environmental exposures. The features, advantages, and limitations of the Cultex® Radial Flow System, the Vitrocell® Continuous Flow and Cloud Alpha systems, and the Scireq® ExpoCube® have been previously reviewed [16–19]. Their major strengths lie in the precise dosimetry, modularity for different exposures, and accommodation of standard ALI culture inserts. Sampling and quality assessment of the airborne stimuli can be accomplished via proprietary equipment with extensive data validating the reproducibility of airborne mixtures and efficiency of particle deposition on ALI culture inserts. Some systems utilize advanced deposition attraction techniques such as thermophoresis in the Scireq® ExpoCube® or electrostatic deposition in the Cultex® Radial Flow System, further improving deposition efficiency and reproducibility of applied doses. The Cultex®, Vitrocell®, and Scireq® systems are modular in the generation of different airborne stimuli, including gases, dry powders, and liquid aerosols. Each system can accommodate different brands of ALI culture inserts, including Transwell™, Falcon®, or Millicell® inserts. Some systems can accommodate different sizes of inserts, such as the Cultex® Radial Flow System with 6-, 12-, and 24-well modules, or the Vitrocell® Continuous Flow and Cloud Alpha systems with modules for up to 96-well plate Transwell™ inserts.

Although commercial *in vitro* exposure systems provide confidence in the reproducibility and accuracy of exposures, they are inaccessible to research groups in which funding cannot support procurement and operation of costly and complex systems. The advantageous features of commercial systems including modularity of ALI insert size, airborne stimulus generation, and sampling quantification are often supplied individually, requiring the purchase of several modules to form a complete system [17, 18, 30]. The multi-component system must then be assembled, operated, and maintained by experienced individuals who may require training. Commercial exposure systems also require adequate space and infrastructure that is not available in all laboratories. Alternative in-house, bespoke exposure systems exist, but are often either limited to low-throughput or have not been validated for exposure uniformity and/or reproducibility [10, 20, 21]. The inadequacy of a simple chamber-style exposure system for applying uniform exposures to well plates was demonstrated (**Figure 1**). To ensure homogenous gas concentration or particle distribution in a chamber system, the air within the chamber must then be mixed and subsequently stagnated for a given duration [19]. In the case of combusted stimuli, stagnation ages smoke which increases cytotoxicity and changes the physical composition of particulate sizes [18, 31]. Similarly, the size of liquid aerosol droplets changes with stagnation due to evaporation and merging of droplets upon collision [32]. Furthermore, for airborne stimuli that are expensive or difficult to generate, the non-direct nature of chamber-style exposures results in low deposition efficiency, with most of the stimulus being deposited on surfaces rather than the cells in culture or remaining suspended in the air of the chamber [19]. The disadvantages of a simple chamber-style exposure system suggest that there is an unmet need for *in vitro* exposure systems that are more complex and adaptable while remaining accessible and, most importantly, reproducible.

Research groups interested in airborne exposure science may benefit from an inexpensive, simple, and disposable manifold for applying uniform airborne exposures across standard well plates. Using our novel manifold systems, the low coefficients of variance in the velocity, pressure, and shear stress distributions observed in the manifold fluid dynamics simulations suggest that gases are effectively divided among well plates (**Figure 3**). The low variance in the deposition of nebulized FITC-dextran demonstrated the ability of the manifolds to uniformly distribute airborne particles as heavy as water aerosols (**Figure 4**). Particles with lower density than water would also be expected to distribute uniformly. We also demonstrated the ability of the manifolds to apply distinct, low variance doses of cannabis concentrate vapour with different commercially available vaporizers (**Figure 5**). These results validate the manifolds for applying reproducible gas and aerosol exposures to well plates with potential applications in cannabis concentrate respiratory exposure research.

Although missing the precise dosimetry and quality control available in commercial systems, open-source manifolds provide an accessible avenue for performing exposure research. Their disposability allows for experimentation with hazardous agents without contamination and sterilization of a complex exposure system. The manifolds also may be re-used in applications where the exposure stimulus can be cleaned from the manifold airways. The method of input perfusion can be variable including hand-driven syringe puffs, smoking or puff-generating machines, continuous-flow nebulizers, or gas cylinder regulators among other potential inputs. Laboratories with access to stereolithography (SLA) 3D printers may fabricate the manifolds on demand. SLA printing is a reliable and affordable method for rapid manufacturing of complex 3D constructs with resolutions as low as 50 µm in the XY plane and 10 µm in the Z axis using the current highest quality SLA printers such as Formlabs Form4. Manufacturing via SLA printing does not require extensive training or experience to reproducibly fabricate parts, with a range of widely accessible brands and models having diverse price points. For custom applications, rapid prototyping and optimization of novel tools is possible at lower costs using 3D printing compared to conventional manufacturing technologies such as injection molding [10, 33]. With respect to the airborne exposure manifolds, few manufacturing technologies can fabricate thin hollow channels with the precision and speed SLA printing provides, with a minimum print time of 3 h using a Formlabs Form4 printer. Each manifold consumes 75 mL and 100 mL of resin while costing $15 and $20 (CAD) per manifold for the 6- and 24-well formats, respectively, using Formlabs Clear V4 resin. Laboratories without direct access to SLA 3D printers may outsource to third party 3D printing services such as Protolabs Network (hubs.com).

The manifolds were designed to accommodate 6- and 24-well plates from Corning®, Eppendorf, and Falcon®. These brands are standard for cell and tissue culture, and they are suitable for standard brands of ALI permeable membrane inserts including Transwell™, Falcon®, and Millicell®. ALI cultures are important models for *in vitro* exposure research because they enable direct airborne exposure to cells, which better reproduces the *in situ* exposure dynamics compared to aqueous applications such as smoke conditioned media or particle suspensions. In the context of lung exposure research, airway epithelial cells differentiate when cultured in ALI conditions, adopting morphology, histology, and gene expression that is more comparable to *in vivo* airway epithelial cells than submerged monolayer cultures [13, 14].

Tobacco smoke exposure has been shown to induce functional changes in human airway epithelial cells [11, 12, 23, 34]. Reductions in TEER are reported in ALI-differentiated Calu-3 cells when exposed to whole tobacco smoke, without apparent changes in morphology or evidence of significant cell death [35]. Tobacco smoke-induced increases in production of IL-6 and IL-8 are reported in Calu-3, MM-39, and NCI-H292 cell lines, and in primary human bronchial epithelial cells [36, 37]. Exposure to soluble constituents of tobacco smoke has been shown to attenuate the activity of ion channels such as CFTR in Calu-3 epithelial cells [12]. To validate the manifolds for investigating the effects of airborne exposures on ALI cell cultures, we applied whole tobacco smoke to ALI cultures of Calu-3 bronchial epithelial cells and measured functional changes that were consistent with similar studies using commercial exposure systems [17, 35]. Smoke exposure had reduced the TEER by nearly 50% of the pre-exposure measurements and increased production of IL-6 and IL-8 without significant increase in released levels of LDH, indicating that the measured changes were functional effects of smoke exposure rather than being consequences of cell death (**Figure 6)**. We also observed dose-dependent attenuation of cell membrane potential with tobacco smoke exposure (**Figure 7**), demonstrating the feasibility of the manifolds for performing concentration-response experiments.

In summary, the manifolds presented herein provide an open-source alternative for exploring *in vitro* exposure science. We validated their utility in 6- and 24-well plates using nebulized aerosols, cannabis vapour, and tobacco smoke, demonstrating uniform aerosol distribution across standard well plates, application of precise vapour doses, and induction of functional changes in ALI cell cultures. Applications may be extended to pollutants, allergens, pathogen-laden aerosols, drug aerosols, cannabis smoke, or wildfire smoke.

## Acknowledgements

The authors would like to acknowledge Tracey Campbell and Nikki Henriquez for their mass spectrometry quantification of CBD.

## Take home message

Open-source 3D printed manifolds enable accessibility to exposure science infrastructure for medium throughput vaping, nebulizing, or combustion experiments with human airway epithelial cells grown under air-liquid interface culture conditions.

---

**Supplementary Figure 1.**
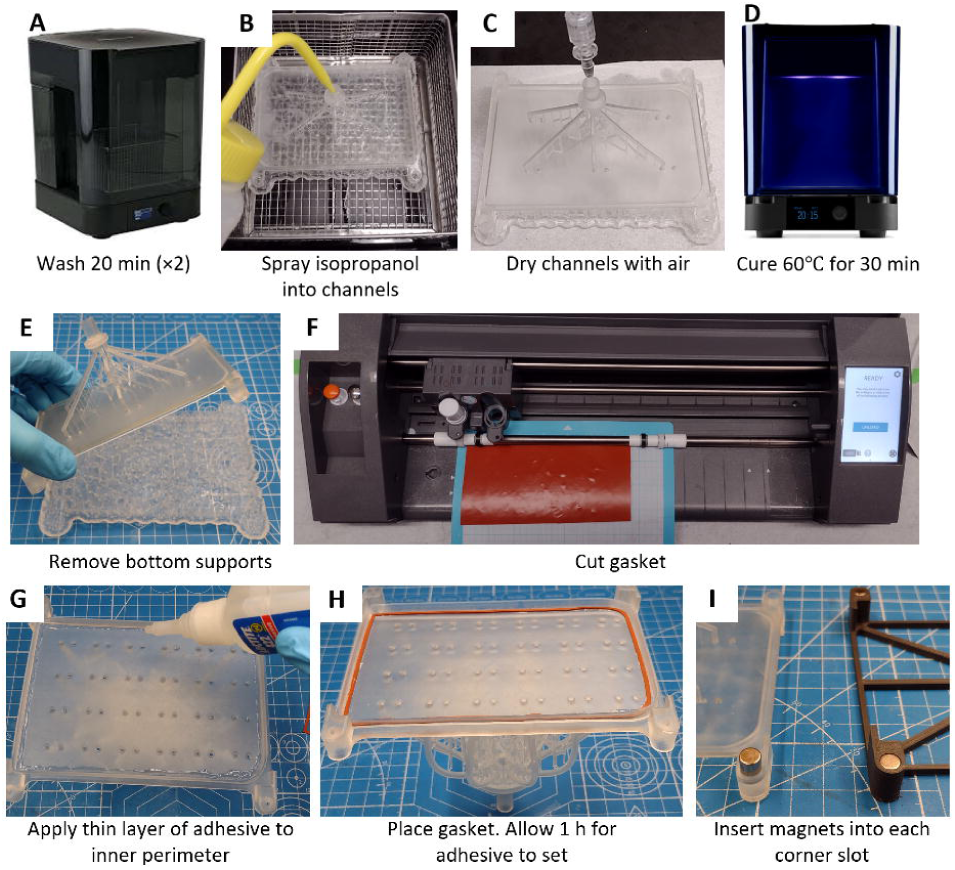
Manifold post-print processing illustrated. **A)** Immediately after removing the manifolds from the printer, they were washed twice in 99% isopropanol (Wisent Inc., product No. 609-400-GL) for 20 min using an automated washing machine (Form Wash, Formlabs, product No. FH-WA-01). **B)** Between washes, a wash bottle (VWR, product No. 10111-954) was used to spray isopropanol into the manifold chamber and channels to ensure all channels were free of uncured resin. **C)** The channels were dried using compressed air before **D)** curing the manifold at 60 for 30 min (Form Cure, Formlabs, product No. FH-CU-01). **E)** All supports were left attached during curing, after which all supports except those attached to the channels were removed. **F-H)** Gaskets were cut from silicone rubber sheets (McMaster Carr, product No. 1460N21) using a cutting machine (Silhouette Cameo 3) and glued to the manifold using instant adhesive (Loctite 422, product No. 233927). The base was fabricated using a fused deposition 3D printer (Ultimaker S3) with PLA filament (Ultimaker, product No. 1609). **I)** Magnets of dimension 6.35×9.52 mm (K&J Magnetics, product No. D46) were inserted into each of the four corner slots of both the manifold and base.

**Supplementary Figure 2.**
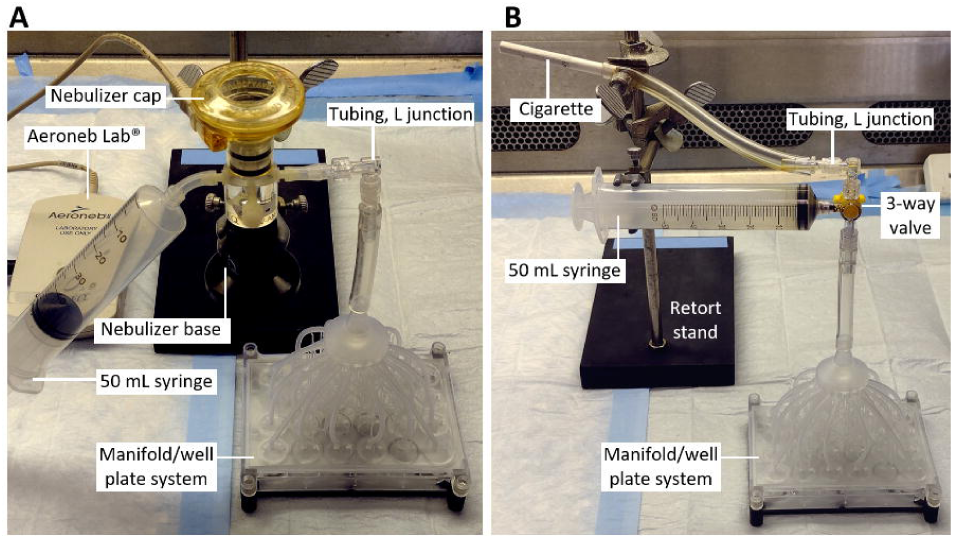
Experimental setups for A) nebulized aerosol and B) tobacco smoke perfusion into the manifolds.

